# LPD-3 as a megaprotein brake for aging and insulin-mTOR signaling in *C. elegans*

**DOI:** 10.1101/2023.02.14.528431

**Authors:** Taruna Pandey, Bingying Wang, Changnan Wang, Jenny Zu, Huichao Deng, Kang Shen, Goncalo Dias do Vale, Jeffrey G. McDonald, Dengke K. Ma

## Abstract

Insulin-mTOR signaling drives anabolic growth during organismal development, while its late-life dysregulation may detrimentally contribute to aging and limit lifespans. Age-related regulatory mechanisms and functional consequences of insulin-mTOR remain incompletely understood. Here we identify LPD-3 as a megaprotein that orchestrates the tempo of insulin-mTOR signaling during *C. elegans* aging. We find that an agonist insulin INS-7 is drastically over-produced in early life and shortens lifespan in *lpd-3* mutants, a *C. elegans* model of human Alkuraya-Kučinskas syndrome. LPD-3 forms a bridge-like tunnel megaprotein to facilitate phospholipid trafficking to plasma membranes. Lipidomic profiling reveals increased abundance of hexaceramide species in *lpd-3* mutants, accompanied by up-regulation of hexaceramide biosynthetic enzymes, including HYL-1 (Homolog of Yeast Longevity). Reducing HYL-1 activity decreases INS-7 levels and rescues the lifespan of *lpd-3* mutants through insulin receptor/DAF-2 and mTOR/LET-363. LPD-3 antagonizes SINH-1, a key mTORC2 component, and decreases expression with age in wild type animals. We propose that LPD-3 acts as a megaprotein brake for aging and its age-dependent decline restricts lifespan through the sphingolipid-hexaceramide and insulin-mTOR pathways.

---

What causes aging remains a contentious unresolved question. Major theories of aging, including the molecular damage and free radical theory, the antagonistic pleiotropy and hyperfunction theory of aging, have stimulated many studies to test predictions from each^1–8^. A new era in aging research was initiated by the discovery of exceptionally long-lived mutants in *C. elegans*, a genetically tractable model organism particularly suited for aging studies^9–12^. Reduced abundance or activity in components of the insulin pathway (the insulin receptor DAF-2 or PI-3 kinase AGE-1) extends longevity through mechanisms that have been extensively studied, and involve, at least in part, activation of geroprotective transcription factors (DAF-16, HSF-1 and SKN-1)^13–15^. While such loss-of-function models have been highly valuable and support critical roles of insulin-mTOR in aging from diverse organisms, whether and how insulin-mTOR hyperfunction may actively drive aging and age-associated pathological phenotypes remain underexplored. Key regulators that can prevent insulin-mTOR hyperfunction and orchestrate the tempo of aging remain unidentified.

In recent studies, we identified *lpd-3* in a mutagenesis screen and discovered that the 452 kDa megaprotein LPD-3 acts as a bridge-like tunnel protein (BLTP) with evolutionarily conserved roles in phospholipid trafficking from the endoplasmic reticulum (ER) to plasma membranes (PM) (Fig. 1a)^16^. Mutations in *BLTP1*, the human orthologue of *lpd-3*, cause a rare autosomal recessive genetic disorder, Alkuraya-Kucinskas syndrome (AKS)^17–19^. Since most AKS patients die prematurely of unknown etiology, we sought to determine the lifespan of *lpd-3* mutants and explore how LPD-3 deficiency causes pathological phenotypes in *C. elegans*. We generated two backcrossed strains carrying a deletion allele *ok2138* or CRISPR-generated stop-codon allele *wy1772*, respectively (Fig. 1b). We found that both strains showed strikingly shortened lifespans at three common temperature cultivation conditions tested (Fig. 1c-e). In addition, we found that *lpd-3* mutants exhibited various common aging phenotypes, including age pigment and reduced behavioral locomotion that occurred earlier than in the wild type (Extended Data Fig. 1).

**Fig. 1.**
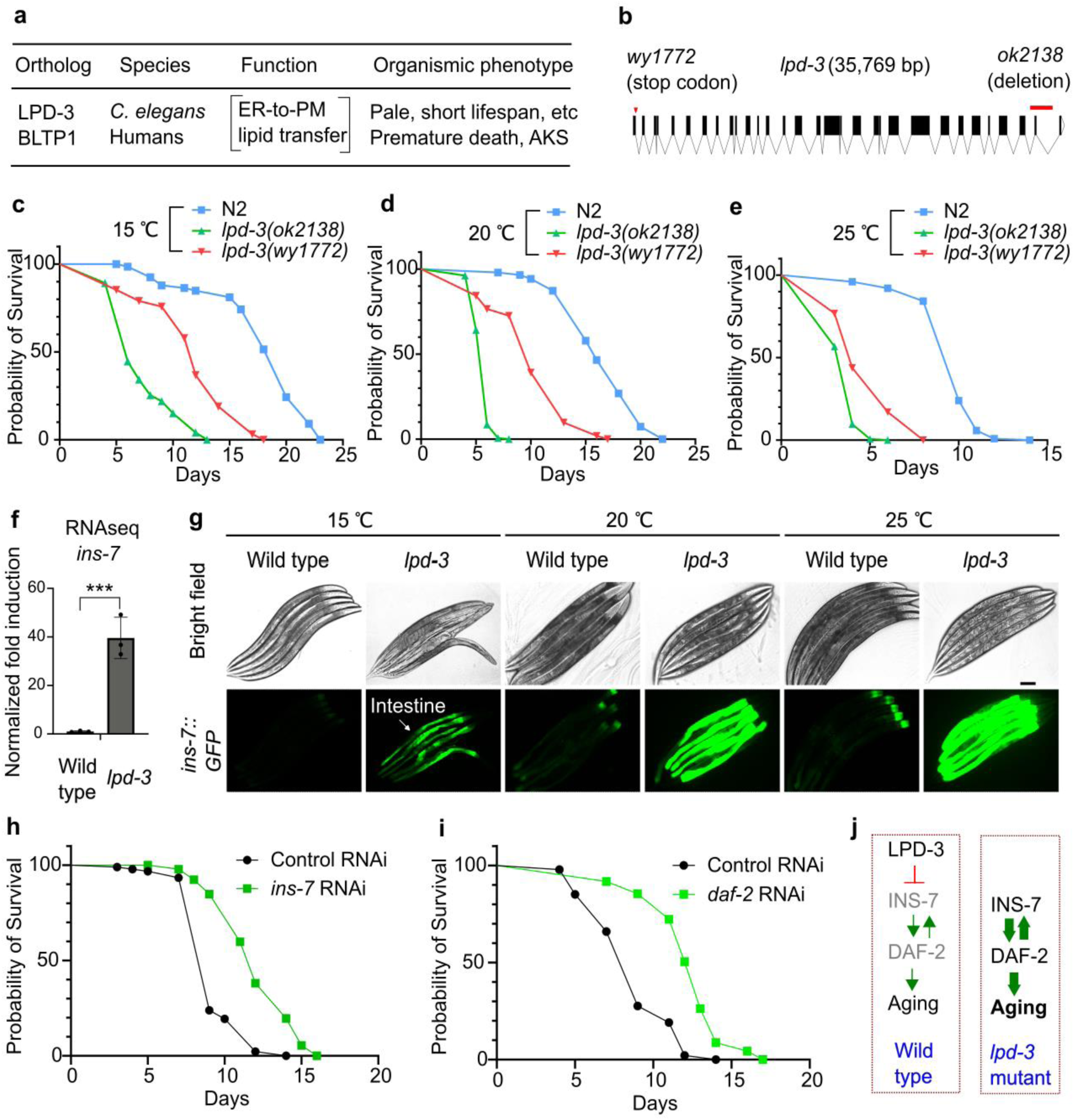
Hyper-activation of *ins-7* causes shortened lifespan in *lpd-3* mutants. **a**, Cellular function and organismic phenotypes for LPD-3 and the human homologue BLTP1. **b**, Gene diagram showing two alleles of *lpd-3* examined for lifespan phenotypes. **c**, Lifespan analysis of wild type versus *lpd-3* mutants with two different alleles (*ok2138* and *wy1772*) grown at 15 °C. **d**, Lifespan analysis of wild type versus *lpd-3* mutants with two different alleles (*ok2138* and *wy1772*) grown at 20 °C. **e**, Lifespan analysis of wild type versus *lpd-3* mutants with two different alleles (*ok2138* and *wy1772*) grown at 25 °C. **f**, RNA-seq results showing increased *ins-7* expression in *lpd-3* mutants. Values are means ± S.D. *** indicates *P* < 0.001 (N = 3 biological replicates). **g**, Representative bright-field and epifluorescence images showing drastically increased *ins-7*p::*ins-7*::GFP abundance in *lpd-3* mutants. Scale bar, 100 µm. **h**, Lifespan analysis of *lpd-3* mutants with control or *ins-7* RNAi at 20 °C, showing shortened lifespan rescued (*P* < 0.0001, log-rank test). **i**, Lifespan analysis of *lpd-3* mutants with control or *daf-2* RNAi at 20 °C, showing shortened lifespan rescued (*P* < 0.0001, log-rank test). **j**, Schematic diagrams illustrating a model of how LPD-3 regulates aging via INS-7 and DAF-2 in wild type and *lpd-3* mutants.

The drastically shortened lifespan of *lpd-3* mutants prompted us to identify the underlying mechanisms. By analyzing RNA-seq datasets for *lpd-3* mutants compared with wild type^16^, we discovered that LPD-3 deficiency led to a nearly 40-fold increase in the expression level of *ins-7*, which represents one of the most highly *lpd-3* up-regulated genes and encodes an agonist insulin with previously reported effects on aging and lifespan in *C. elegans*^20–22^ (Fig. 1f). Expression of other insulin-encoding genes was much less regulated or largely unaltered (Extended Data Fig. 2a, b). To determine the spatiotemporal site of *ins-7* regulation, we crossed an integrated *ins-7*p::INS-7::GFP translational reporter strain^23^ with the *lpd-3* deletion mutant. In the wild type, *ins-7*p::INS-7::GFP is expressed at baseline levels that are detectable only in the anterior part of intestine at 15 °C starting from young adult stages (24 hrs post-L4), and increased at 20 °C and 25 °C (Fig. 1g). By contrast, *ins-7*p::INS-7::GFP in *lpd-3* mutants showed drastically increased INS-7::GFP abundance and fluorescence throughout the intestine, and particularly under higher temperatures (20 or 25 °C). We showed previously that *lpd-3* mutants are defective in ER-to-PM phospholipid trafficking, and exhibit sensitivity to thermal stress that can be normalized by Lecithin^16^. We confirmed in this study that Lecithin supplementation in the culture medium rescued both the shortened lifespan and *ins-7* up-regulation phenotypes (Extended Data Fig. 2c, d).

To determine the causal role of *ins-7* in the shortened lifespan phenotype of *lpd-3* mutants, we used RNAi to knock down *ins-7* expression in *lpd-3* mutants. We found that treatment with RNAi against *ins-7* led to a markedly extended lifespan of *lpd-3* mutants, comparable to that in the wild type (Fig. 1h). We also observed such rescue of shortened lifespan phenotype of *lpd-3* mutants by RNAi against *daf-2* (Fig. 1i), which encodes the only known receptor of insulin in *C. elegans*. We note that control RNAi also led to a slight increase of lifespan in *lpd-3* mutants (Fig. 1d, h), likely reflecting different lipid compositions of bacterial strains between OP50 and HT115 strains^24, 25^ used in routine *C. elegans* culture and RNAi experiments, respectively. Reducing *ins-7* expression has been previously shown to increase the lifespan of wild-type animals, implicating INS-7’s role in tissue entrainment by feedback regulation in *C. elegans*^20–22^. Our results support these early findings and reveal that overproduction of INS-7 is causally responsible for shortening lifespan in *lpd-3* mutants, in a manner that requires the sole insulin receptor DAF-2 (Fig. 1j).

We next determined the regulatory mechanisms leading to *ins-7* up-regulation and shortened lifespan in *lpd-3* mutants. Taking advantage of the drastically up-regulated *ins-7*p:: INS-7::GFP reporter in *lpd-3* mutants as a live fluorescent readout, we performed targeted RNAi screens to identify genes required for *ins-7*p:: INS-7::GFP up-regulation by *lpd-3*. We performed RNAi for those genes with reported adequate expression in the intestine (transcript per million, TPM > 2) and encoding mediators of major signal transduction pathways known to be involved in insulin signaling (the Ras/MAPK, phospholipase and PI3 kinase/mTORC1/mTORC2 pathways)^26–28^ (Extended Data Fig. 3). As positive controls, we verified strong effects of *ins-7* or *daf-2* RNAi in suppressing *ins-7*p::INS-7::GFP expression in *lpd-3* mutants (Fig. 2a, b). From such screens, we found that RNAi against several genes encoding components of the mTORC2 complex in *C. elegans*, including *let-363*, *rict-1*, and *sinh-1*, led to a marked reduction of *ins-7*p::INS-7::GFP in *lpd-3* mutants (Fig. 2a, b). RNAi against genes involved in the mTORC1 complex, except *let-363* shared by mTORC1 and mTORC2, exhibited weaker effects (Fig. 2a).

**Fig. 2.**
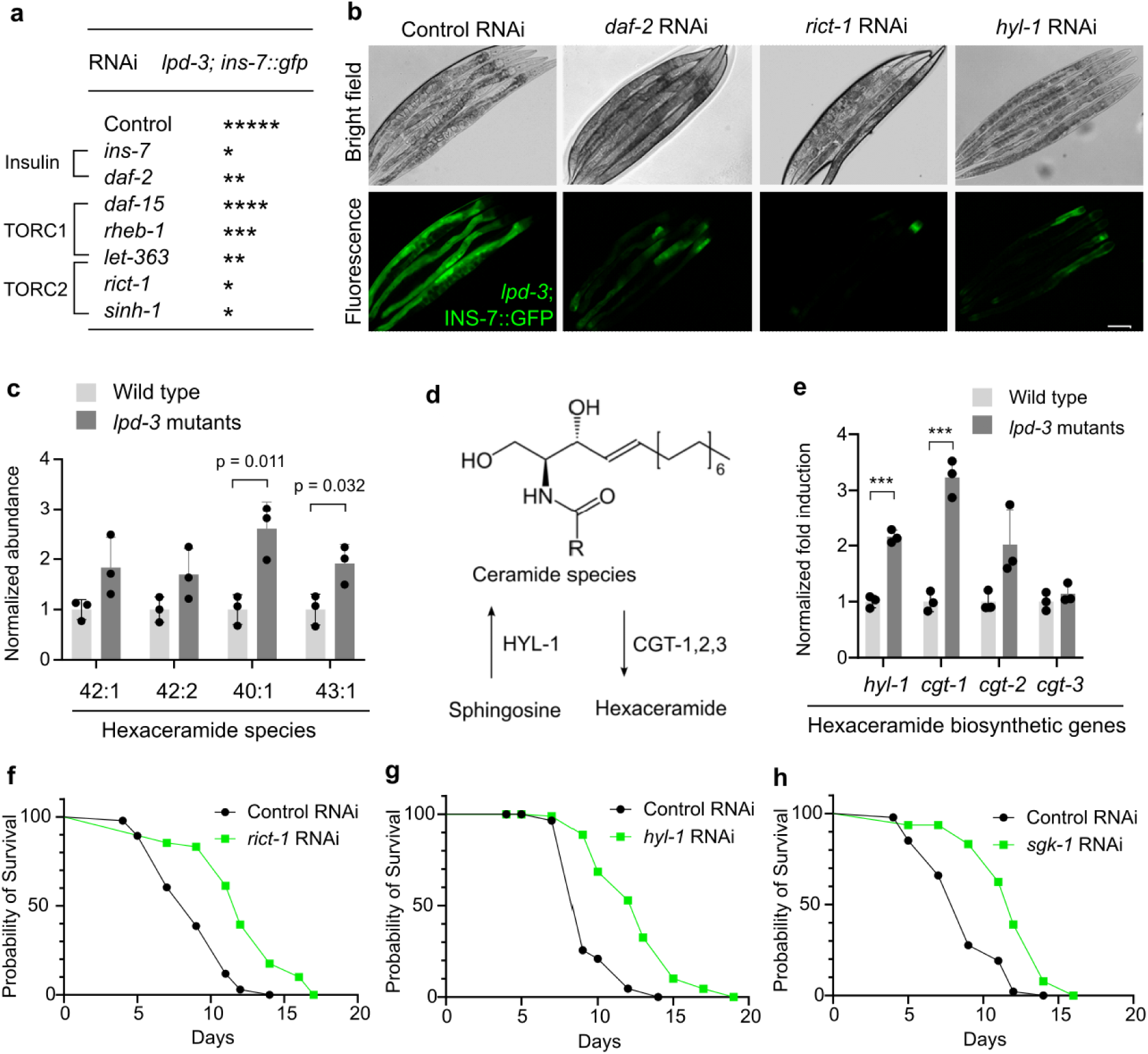
LPD-3 regulates INS-7 via the sphingolipid-ceramide-mTORC2 axis. **a**, Table summarizing effects of RNAi against genes in the insulin/mTORC1/mTORC2 pathway on *ins-7*p::*ins-7*::GFP levels in *lpd-3* mutants. * indicates GFP intensity observed under microscope. **b**, Representative bright-field and epifluorescence images showing *ins-7*p::*ins-7*::GFP up-regulation in *lpd-3* mutants can be suppressed by RNAi against *daf-2*, *rict-1* or *hyl-1*. Scale bar, 100 µm. **c**, Lipidomic quantification of hexaceramide species in wild type and *lpd-3* mutants. Values are means ± S.D. (N = 3 biological replicates). **d**, Schematic showing biosynthetic pathways of hexaceramide, including sphingosine conversion to ceramide by HYL-1 and ceramide to hexaceramide by CGT-1/2/3. **e**, RNA-seq results showing increased *hyl-1* and *cgt-1/2/3* expression in *lpd-3* mutants. Values are means ± S.D. \*\*\**P* < 0.001 (N = 3 biological replicates). **f**, Lifespan analysis of *lpd-3* mutants with control or *rict-1* RNAi at 20 °C, showing rescued lifespan (*P* < 0.0001, log-rank test). **g**, Lifespan analysis of *lpd-3* mutants with control or *hyl-1* RNAi at 20 °C, showing rescued lifespan (*P*< 0.0001, log-rank test). **h**, Lifespan analysis of *lpd-3* mutants with control or *sgk-1* RNAi at 20 °C, showing rescued lifespan (*P* < 0.0001, log-rank test).

In parallel to such RNAi screens, we also performed lipidomic profiling of *lpd-3* mutants compared with wild type. Among all major lipid species examined by LC-MS/MS, including phospholipids, glycerolipids and sphingolipids (Supplementary Table 1), the hexaceramide-type of sphingolipids showed marked up-regulation in *lpd-3* mutants (Fig. 2c). Consistently, RNA-seq results revealed that genes encoding hexaceramide biosynthetic enzymes (Fig. 2d), including HYL-1 and CGT-1/2/3, were markedly up-regulated in *lpd-3* mutants (Fig. 2e). As reduced LPD-3 activity results in impaired ER-to-PM phospholipid trafficking, a lipid homeostatic mechanism is likely triggered to up-regulate sphingolipids in *lpd-3* mutants^16^. Given reported sphingolipid modulation of mTOR signaling and longevity^29–31^, we next explored the link of hexaceramide biosynthetic enzyme HYL-1 to *ins-7*. We found that *hyl-1* RNAi led to strong suppression of *ins-7*p::INS-7::GFP in *lpd-3* mutants (Fig. 2b). Functionally, lifespan analysis revealed that RNAi against *rict-1* or *hyl-1* led to the marked rescue of the shortened lifespan in *lpd-3* mutants (Fig. 2f, g). RNAi against *sgk-1*, which encodes a major kinase^32–34^ mediating effects of mTORC2, showed similar rescuing effects (Fig. 2h). Taken together, these results indicate that the sphingolipid-mTOR pathway drives *ins-7* over-expression, leading to the shortened lifespan in *lpd-3* mutants.

We next explored cellular mechanisms by which LPD-3 regulates mTORC2. The mTORC2 complex in *C. elegans* consists of the core subunit LET-363 shared by mTORC1, the mTORC2-specific subunits RICT-1 and SINH-1 (Fig. 3a). As the mammalian homologs of SINH-1 bind to PIP3 at the PM and mediate signaling from insulin receptors to the mTORC2 complex^35–, 37^, we tagged endogenous SINH-1 with GFP by CRISPR (Fig. 3b) and monitored how LPD-3 may affect the intracellular abundance and localization of SINH-1::GFP. In the wild type, we did not observe apparent SINH-1::GFP fluorescence throughout animals at the young adult stage (24 hrs post L4). By contrast, RNAi against *lpd-3* or its mutations led to the marked increase of SINH-1::GFP fluorescence signals that were particularly enriched at loci in close proximity to PM in the intestine (Fig. 3c, d). As expected, RNAi against *lpd-3* also effectively decreased the abundance of LPD-3 endogenously labeled with GFP by CRISPR (Fig. 3e, and see below). Since loss of LPD-3 causes PIP2/3 to accumulate in the ER membrane near the ER-PM junction^16^, increased SINH-1::GFP in *lpd-3* mutants likely reflects its ectopic intracellular activity. In addition to the *ins-7* phenotype (Fig. 2), we found that *lpd-3* mutants exhibited morphologically spherical mitochondria with decreased branch lengths, whereas RNAi against genes encoding the mitochondrial fission machinery (DRP-1) or insulin-mTOR pathway components (INS-7, DAF-2, SINH-1, SGK-1) suppressed such phenotype (Fig. 3f, g). These results indicate that LPD-3 normally functions to restrict SINH-1 accumulation and thereby antagonize ectopic mTOR activity.

**Fig. 3.**
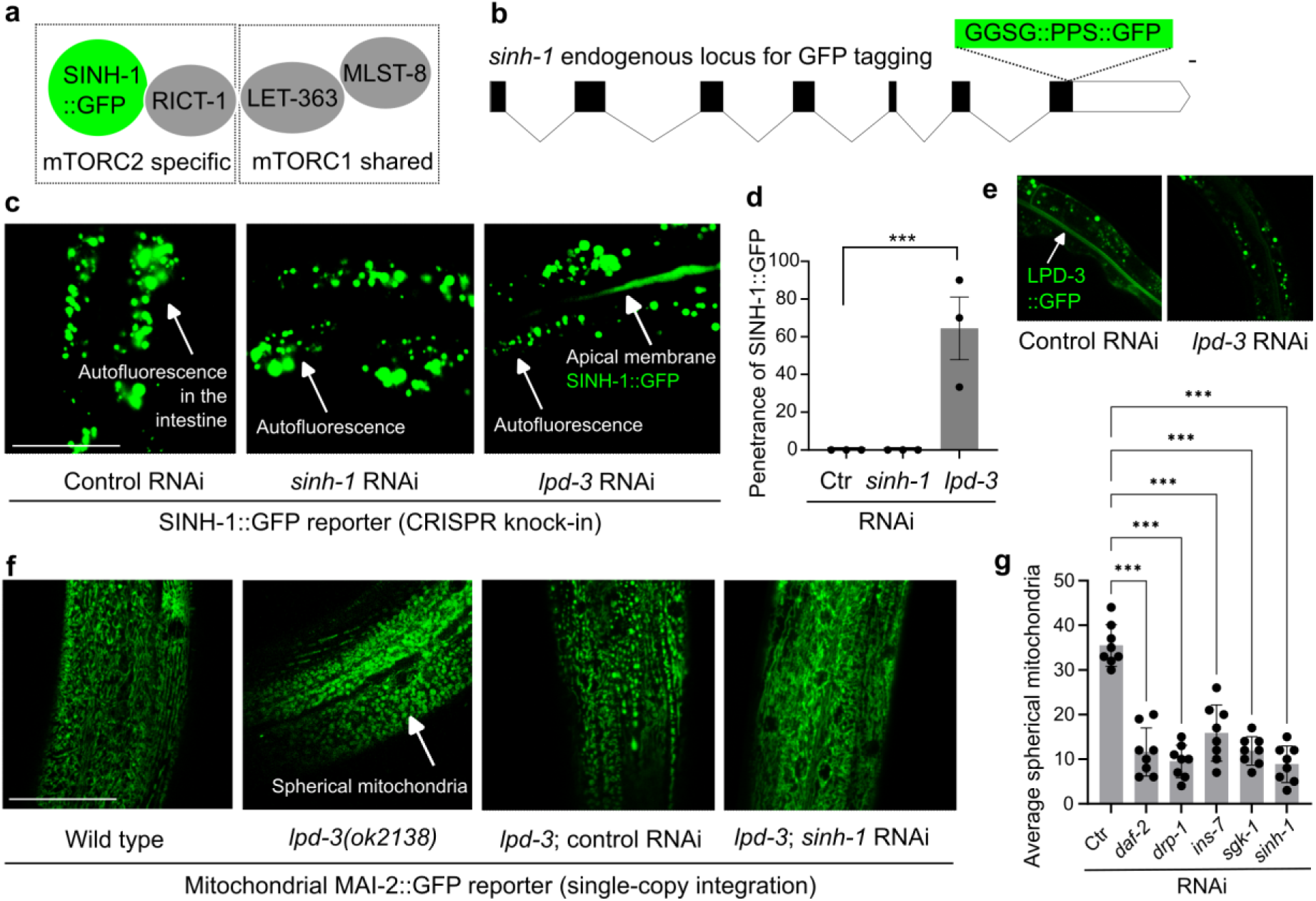
Loss of LPD-3 dysregulates SINH-1/mTORC2 and mitochondria. **a**, Schematic showing key components of the mTORC2 complex in the SINH-1::GFP tagged *C. elegans.* **b**, Schematic gene structure of *sinh-1* showing the CRISPR-mediated knock-in allele encoding the endogenous SINH-1 tagged with a linker (GGGS), PreScission Protease site (PPS) and GFP. Scale bar: 100 bp. **c**, Representative confocal fluorescence images showing undetectable SINH-1::GFP signals in animals treated with control or *sinh-1* RNAi but increased SINH-1::GFP enrichment along the apical intestinal membrane in animals treated with *lpd-3* RNAi (non-specific gut autofluorescence not affected by *sinh-1* or *lpd-3* RNAi also indicated). **d**, Quantification of the penetrance of SINH-1::GFP enrichment at the apical intestinal membrane in animals with indicated RNAi treatment. Values are means ± S.E.M. \*\*\**P* < 0.001 (N = 3 independent experiments, n > 10 in each trial). **e**, Representative confocal fluorescence images showing decreased LPD-3::GFP by *lpd-3* RNAi. **f**, Representative confocal fluorescence images showing increased spherical MAI-2::GFP marked mitochondria in *lpd-3* mutants (day 1) that can be normalized by *sinh-1* but not control RNAi. **g**, Quantification showing rescued mitochondrial morphological defects in *lpd-3* mutants by RNAi against genes encoding the mitochondrial fission machinery (*drp-1*) or insulin-mTOR pathway components (e.g. *sinh-1*). Values are means ± S.E.M. \*\*\**P* < 0.001 (N = 8 biological replicates).

Are endogenous LPD-3 abundance and activity subject to regulation by normal aging in wild type animals? To address this question, we tagged endogenous LPD-3 with seven copies of the split GFP (GFP11) by CRISPR^38–40^, crossed the allele into a strain expressing GFP10 specifically in the intestine by the *ges-1* promoter (Fig. 4a). We monitored how aging affects the intracellular abundance and localization of LPD-3::GFP based on the complementation of GFP1-10 and 7xGFP11 in the intestine, site of which we showed previously is critical for LPD-3 functions in phospholipid trafficking^16^. Confocal microscopy revealed that LPD-3::GFP is enriched along the apical membrane of the intestine at the L4 and young adult stages, with an age-dependent progressive decline in abundance at the stages of day 3, 5 and 9 (Fig. 4b, c). Functionally, we showed previously that LPD-3 facilitates ER-to-PM trafficking of phospholipids, including phosphatidylcholine and phosphatidylinositol. Using Akt-PH::GFP, which binds to the phosphatidylinositol species PIP2/PIP3, we found that aging caused a progressive decline of Akt-PH::GFP signals at the intestinal apical membrane while *lpd-3* loss-of-function mutations exacerbated such age-dependent decline (Fig. 4d, e). By contrast, *ins-7*p::INS-7::GFP exhibited age-dependent increase in abundance while such age-dependent change was accelerated in *lpd-3* mutants (Fig. 4f, g). These results demonstrate the age-dependent decline in both the abundance and functional activity of endogenous LPD-3 during normal *C. elegans* aging.

**Fig. 4.**
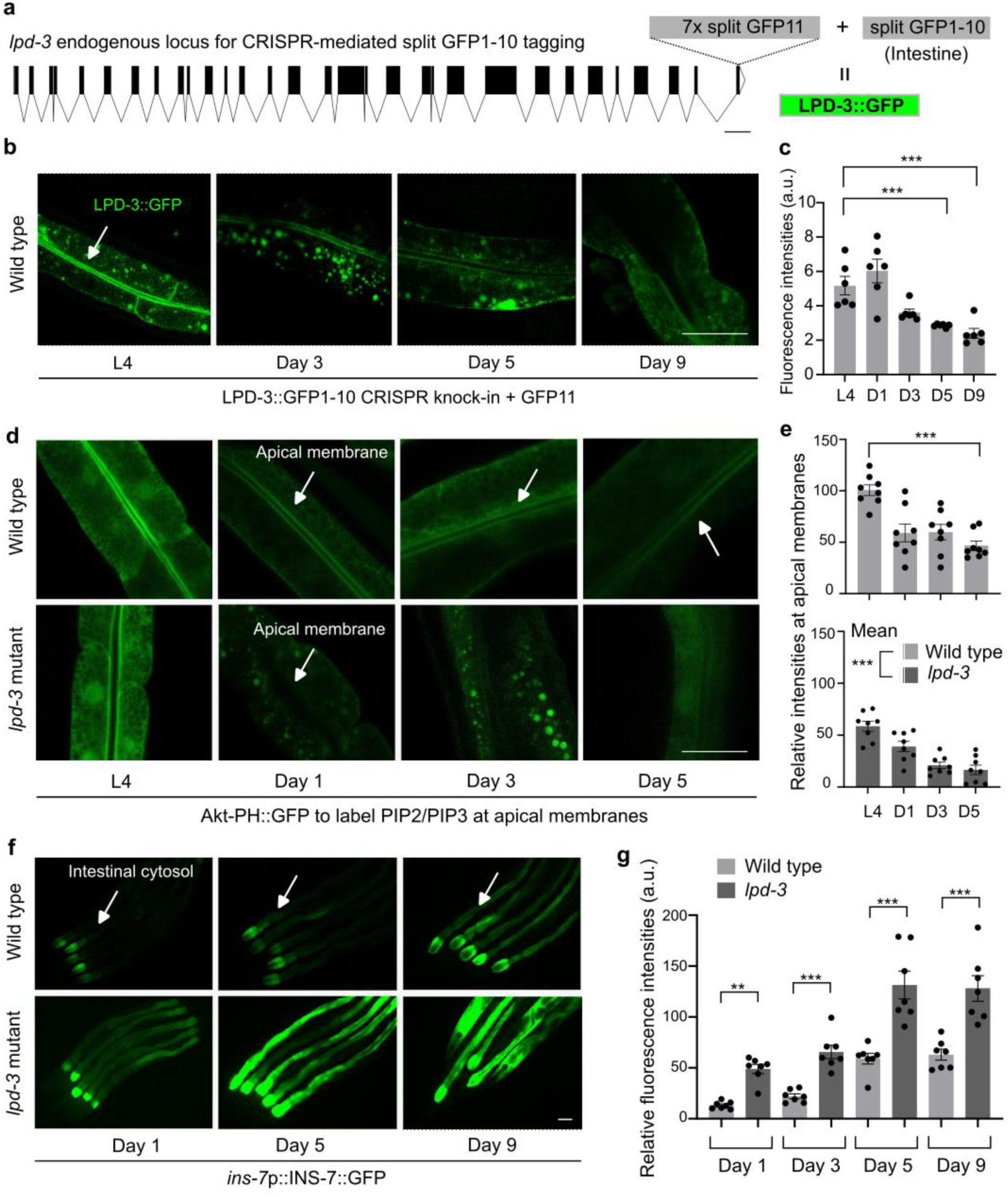
LPD-3 abundance and functions decline with age in wild type animals. **a**, Schematic gene structure of *lpd-3* showing the CRISPR-mediated knock-in allele encoding the endogenous LPD-3 tagged with a split GFP1-10 that is complemented by an intestine-expressed GFP11. Scale bar: 500 bp. **b**, Representative confocal fluorescence images showing age-dependent decrease in the abundance of LPD-3::GFP signals in the intestinal apical membrane of animals at the stages of L4, day 1, 3, 5 and 9 post-L4. **c**, Quantification of the fluorescence intensities of LPD-3::GFP in animals with indicated ages. Values are means ± S.E.M. \*\*\**P* < 0.001 (N = 6 biological replicates). **d**, Representative confocal fluorescence images showing age-dependent decrease in the abundance of Akt-PH::GFP signals that label PIP2/3 in the intestinal apical membrane of animals at the stages of L4, day 1, 3, 5 and 9 post-L4, in both wild type and *lpd-3* mutants. Scale bar: 50 µm. **e**, Quantification of the fluorescence intensities of Akt-PH::GFP at apical intestinal membranes in animals with indicated ages. Values are means ± S.E.M. \*\*\**P* < 0.001 (N = 8 biological replicates in each stage, N = 32 in each genotype). **f**, Representative confocal fluorescence images showing age-dependent increase in the abundance of *ins-7*p::INS-7::GFP signals in animals at the indicated stages in both wild type and *lpd-3* mutants. Scale bar: 50 µm. **e**, Quantification of the fluorescence intensities of *ins-7*p::INS-7::GFP in animals with indicated ages. Values are means ± S.E.M., with ** *P* < 0.01 and \*\*\**P* < 0.001 (N = 7 biological replicates).

Identification and subsequent mechanistic studies of exceptionally long-lived *C. elegans* mutants in the insulin-mTOR pathway have provided crucial insights into how the loss of insulin-mTOR function extends longevity. Activation of key transcription factors including DAF-16, HSF-1, and SKN-1 mediates geroprotective effects in these mutants by promoting somatic maintenance programs that attenuate molecular and cellular damages. Our studies provide a novel and complementary model in which we propose insulin-mTOR hyperfunction can actively drive aging downstream of LPD-3 and the sphingolipid-ceramide pathway (Extended Data Fig. 4). Our findings support the notion that somatic maintenance programs and molecular/cellular damaging agents (e.g. free radicals and/or cell death signals) from dysregulated mitochondria, both of which act downstream of insulin-mTOR signaling, may converge to control the tempo of aging progression. Based on new insights from *C. elegans* models, this view may imply to reconcile and unify two prevalent mechanistic theories of aging (molecular damage vs TOR hyperfunction)^1–8^.

Various sphingolipids have been shown to contribute to organismal aging and age-associated pathologies^41–43^. We identify LPD-3 as a critical megaprotein regulator of the phospholipid-sphingolipid rheostat, dysfunction of which can profoundly affect aging via insulin-mTOR hyperfunction. LPD-3 defines a member of a recently discovered and highly evolutionarily conserved protein family that controls lipid homeostasis in eukaryotic cell membranes^16, 44–46^; it is currently unknown whether LPD-3 homologs may play similar roles in regulating insulin-mTOR and aging in other organisms. Detailed mechanisms linking LPD-3 to sphingolipid regulation, insulin over-production and aging also await further studies. Based on our findings and prior studies supporting emerging roles of sphingolipid/ceramides and conserved insulin-mTOR signaling from many species^30, 42, 43, 47, 48^, we propose that LPD-3 may act as a cellular brake that slows organismal aging by orchestrating its tempo and regulating the sphingolipid-insulin-mTOR axis likely in phylogenetically diverse organisms.

## Methods

### *C. elegans* culture and maintenance

All the *C. elegans* strains used in the current research were maintained in accordance with the standard laboratory procedures unless otherwise stated. Genotypes of strains used are as follows: N2 Bristol strain (wild type), *lpd-3(ok2138), lpd-3(wy1772), yxIs13[ins-7p::ins-7::gfp; unc-122p::dsred], pwIs890[Pvha-6::AKT(PH)::GFP], xmSi[mai-2::GFP(single copy integration)], sinh-1(syb7471[GGSG::PPS::GFP]), lpd-3(wy1770[7xGFP11]); wyEx10609 [ges-1p:: GFP(1-10); odr-1p::GFP; ges-1p:: ERSP::mScarlet::MAPPER]*.

### Lifespan assessment

The lifespan assay was conducted according to the standard protocol^49^. Briefly, wild type N2 strain was grown for two generations without starvation, and then embryos were synchronized through standard sodium hypochlorite treatment. These embryos were grown on OP50-seeded plates or transferred to RNAi-seeded NGM plates. Thereafter, 50-100 late-L4 larvae or young adults (24 hrs post-L4) were transferred to OP50-seeded plates or RNAi-seeded NGM plates supplemented with 50 μM of 5-fluoro-2-deoxyuridine (FUDR, Sigma) to prevent progeny growth. Assay populations were transferred to fresh OP50-seeded plate every 3-4 days. Live/dead/missing worms were scored on alternate days until the last surviving worm. The live worms were detected through touch provoke responses. The lifespan assays were repeated for at least three independent trials.

### Epifluorescence and confocal microscopic imaging

The epifluorescence compound and confocal microscopes (Leica) were used to capture fluorescence images. Animals of various genotypes or treated with different RNAi were randomly picked and treated with 10 mM sodium azide in M9 solution (Sigma-Aldrich), symmetrically aligned on 2% agar pads on slides for imaging. The control and treatment groups were imaged under identical settings and conditions. The age-dependent decline in LPD-3::GFP strains was scored by calculating the mean fluorescence value from Z-stack of confocal planes in each worm at different time points near the apical membrane using the ImageJ software. Mitochondrial morphology was scored as the number of spherical mitochondria in each image counted manually. The penetrance of SINH-1::GFP in strains of different genotypes under different conditions was scored as a percentage of animals that exhibit detectable fluorescence with confocal microscopy.

### Behavioral and age pigment/lipofuscin measurements during aging

The locomotion of worms was measured as previously described^51^. Briefly, the speed (total track length/time) of age-synchronized worms was recorded using WormLab System (MBF Bioscience) based on the midpoint positions of the worms at different time points. Each experiment was repeated three times with more than 10 animals per group. The age pigment/lipofuscin accumulation with age progression was measured as previously described^52, 53^ by using ultraviolet light excitation coupled with detection through a blue/cyan filter and quantified by calculating the mean fluorescence value in each worm using the ImageJ software.

### Lipidomic analysis

Wild type N2 strain and *lpd-3*(*ok2138*) mutants were grown under non-starved conditions for four generations at 20 °C, followed by bleach synchronization and harvest by M9 buffer at young adult stages. Frozen worm pellets (50 µl per sample, three independent biological replicates per genotype) are stored until lipidomic extraction and analysis. All solvents for lipidomic extraction and analysis used were either HPLC or LC/MS grade and purchased from Sigma-Aldrich (St Louis, MO, USA). Splash Lipidomix standards were purchased from Avanti (Alabaster, AL, USA). All lipid extractions were performed in 16×100mm glass tubes with PTFE-lined caps (Fisher Scientific, Pittsburg, PA, USA). Glass Pasteur pipettes and solvent-resistant plasticware pipette tips (Mettler-Toledo, Columbus, OH, USA) were used to minimize leaching of polymers and plasticizers. Samples were transferred to fresh glass tubes, and 1 mL of methanol, 1mL of water and 2mL of methyl tert-butyl ether (mTBE) were added for liquid-liquid extraction. The mixture was vortexed and centrifuged at 2,671 g for 5 min, resulting in two distinct liquid phases. The organic phase (upper phase) was transferred to a fresh tube with a Pasteur pipette and spiked with 20uL of a 1:5 diluted Splash Lipidomix standard mixture. The samples were dried under N_2_ and resuspended in 400 µL of hexane. Lipids were analyzed by LC-MS/MS using a SCIEX QTRAP 6500^+^ (SCIEX, Framingham, MA) equipped with a Shimadzu LC-30AD (Shimadzu, Columbia, MD) high-performance liquid chromatography (HPLC) system and a 150×2.1 mm, 5 µm Supelco Ascentis silica column (Supelco, Bellefonte, PA). Samples were injected at a flow rate of 0.3 ml/min at 2.5% solvent B (methyl tert-butyl ether) and 97.5% Solvent A (hexane). Solvent B was increased to 5% over 3 min and then to 60% over 6 min. Solvent B was decreased to 0% during 30 sec while Solvent C (90:10 (v/v) isopropanol-water) was set at 20% and increased to 40% during the following 11 min. Solvent C is increased to 44% over 6 min and then to 60% over 50 sec. The system was held at 60% solvent C for 1 min prior to re-equilibration at 2.5% of solvent B for 5 min at a 1.2 mL/min flow rate. Solvent D [95:5 (v/v) acetonitrile-water with 10 mM Ammonium acetate] was infused post-column at 0.03 ml/min. Column oven temperature was 25°C. Data was acquired in positive and negative ionization mode using multiple reaction monitoring (MRM). The LC-MS/MS data was analyzed using MultiQuant software (SCIEX). The identified lipid species were normalized to its corresponding internal standard.

### Statistics and reproducibility

All the data in the current manuscript were analyzed using GraphPad Prism 9.2.0 Software (Graphpad, San Diego, CA) and presented as means ± S.E.M. unless otherwise specified, with significance *P* values calculated by unpaired two-sided *t*-tests (comparisons between two groups), one-way or two-way ANOVA (comparisons across more than two groups) and adjusted with Bonferroni’s multiple comparisons tests. Lifespan assay was quantified using Kaplan–Meier lifespan analysis and *P* values were calculated using the log-rank test.

## Acknowledgments

Some strains were provided by Dr. Yun Zhang’s group at Harvard University, Dr. R. E. Navarro’s group at Universidad Nacional Autónoma de México, and by *Caenorhabditis* Genetics Center (CGC), which is funded by NIH Office of Research Infrastructure Programs (P40 OD010440). The work was financially supported by the NIH/NIGMS grant 1R35GM139618, UCSF PBBR New Frontier Research, and UCSF BARI Investigator Award (D.K.M).

## Author contributions

T.P., B.W., C.W., J.Z. and D.K.M. designed, performed and analyzed most of the *C. elegans* experiments, contributed to project conceptualization and wrote the manuscript. G.V. and J.G.M. contributed to lipid analysis. H.D. and K.S. contributed to the CRISPR allele generation and project conceptualization. D.K.M. supervised the project.

## Materials & Correspondence

Correspondence and material requests should be addressed to Dengke K. Ma, Ph.D.

## Figures and Figure legends

**Extended Data Fig. 1.**
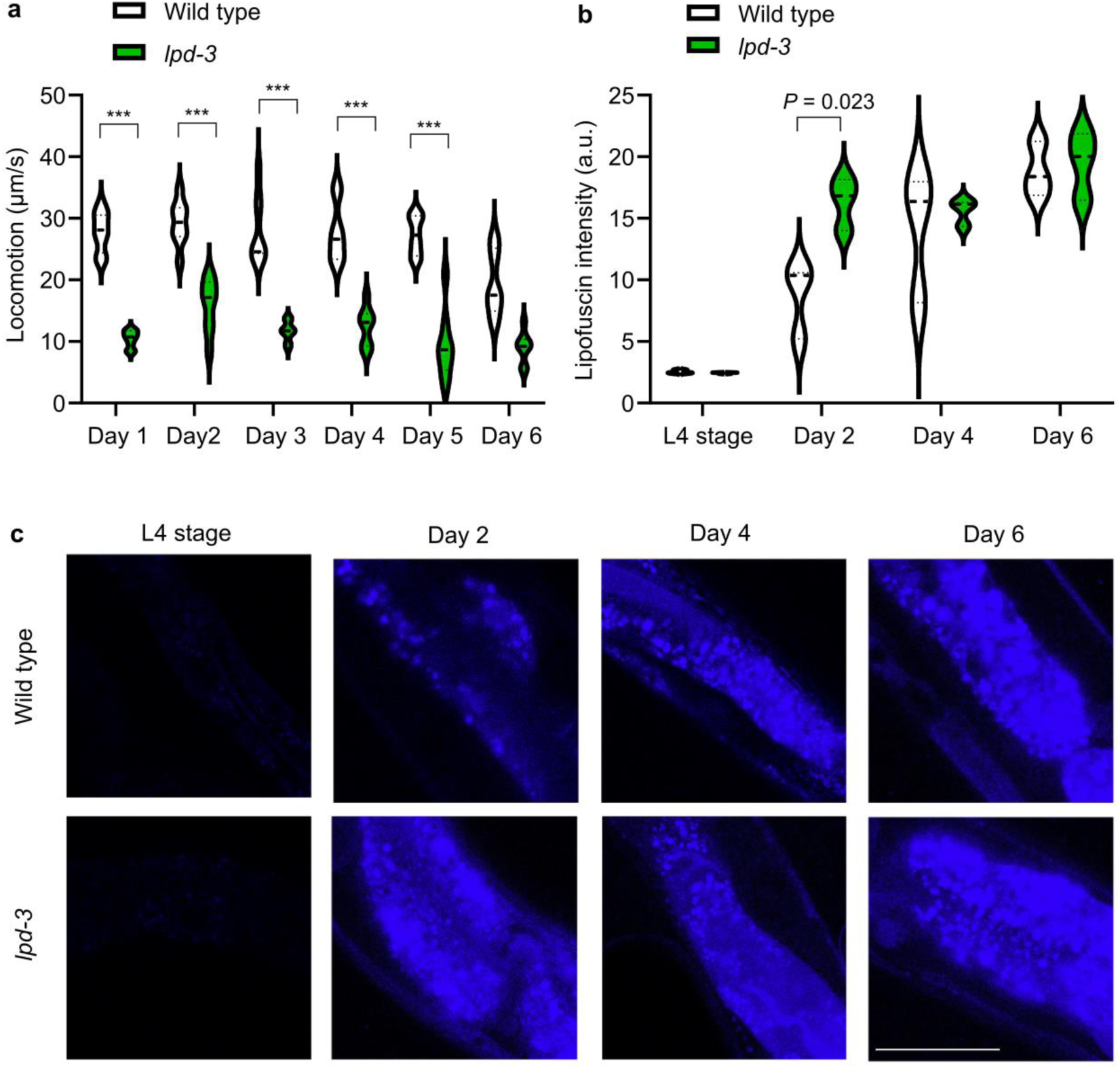
Lipofuscin increase and behavioral decline of *lpd-3* mutants early in aging. **a,** Quantification of locomotion speed for wild type and *lpd-3* mutants showing a rapid decline in *lpd-3* mutants starting at day 1 (24 hrs post-L4 stage). *** indicates *P* < 0.001 (N = 30-50 animals per group). **b,** Lipofuscin/age pigment fluorescence intensities for wild type and *lpd-3* mutants showing a more rapid decline in *lpd-3* mutants at day 2 (48 hrs post-L4 stage). **c**, Representative lipofuscin/age pigment fluorescence images for wild type and *lpd-3* mutants showing a more rapid decline in *lpd-3* mutants at day 2 (48 hrs post-L4 stage). Scale bar, 100 µm.

**Extended Data Fig. 2.**
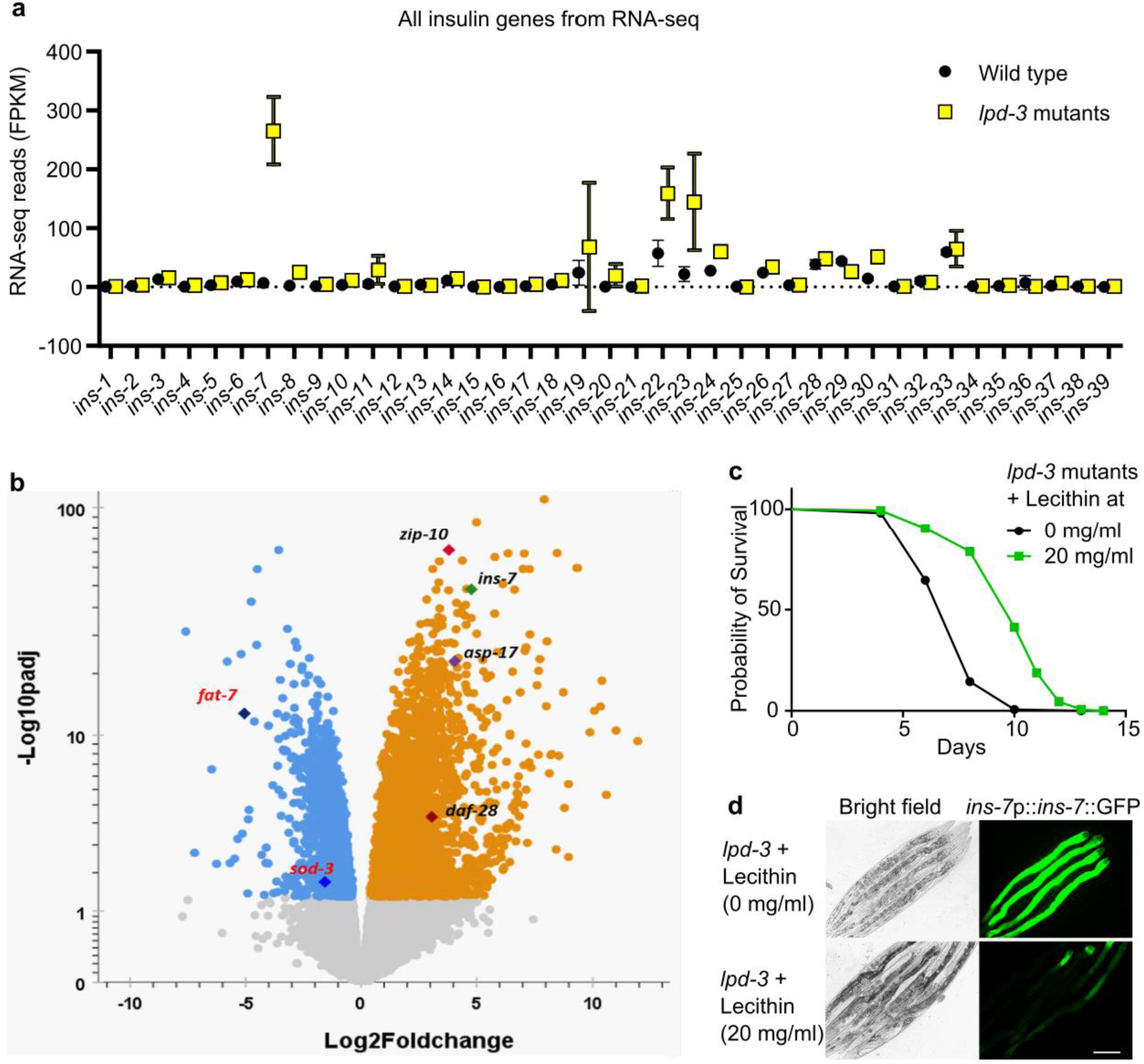
Specific up-regulation of *ins-7* by loss of LPD-3 and rescue by Lecithin. **a**, RNA-seq reveals drastic and specific *ins-7* up-regulation among all *ins* genes examined. Results were derived and analyzed from RNA-seq datasets we published earlier^16^. FPKM, Fragments Per Kilobase of transcript per Million mapped reads. **b**, Volcano plot showing significantly LPD-3 regulated genes, including *fat-7* (downstream of SBP-1 and NHR-49), *sod-3* (downstream of DAF-16) and *zip-10* (downstream of ISY-1), *asp-17* (downstream of ISY-1), and *ins-7*. **c**, Lifespan analysis of *lpd-3* mutants with control or Lecithin treatment at 20 °C (*P* < 0.0001, log-rank test). **d**, Representative bright-field and epifluorescence images showing that *ins-7*p::*ins-7*::GFP up-regulation in *lpd-3* mutants can be suppressed by exogenous Lecithin (20 mg/ml) treatment supplemented in culture media. Scale bar, 100 µm.

**Extended Data Fig. 3.**
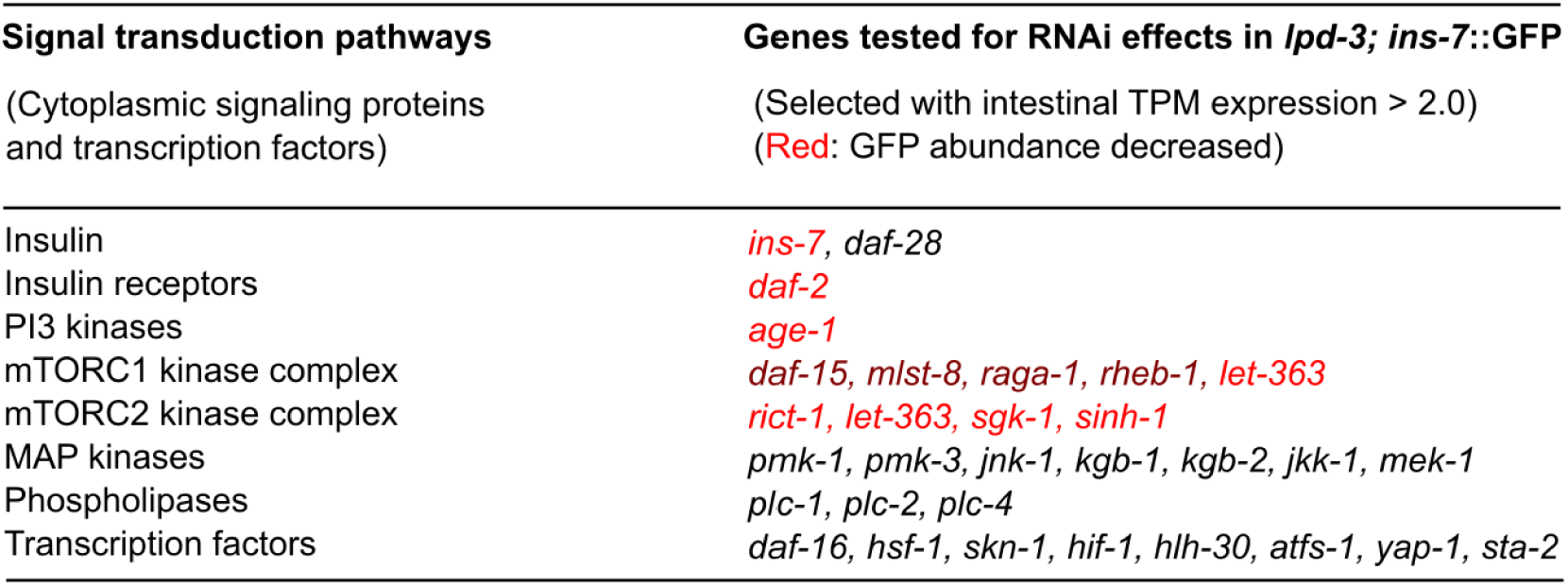
RNAi screens for genes affecting *ins-7* in *lpd-3* mutants. Genes were selected for RNAi testing given their adequate expression values in intestine (TPM or transcript per million > 2.0) and putative roles in mediating insulin-mTOR signaling.

**Extended Data Fig. 4.**
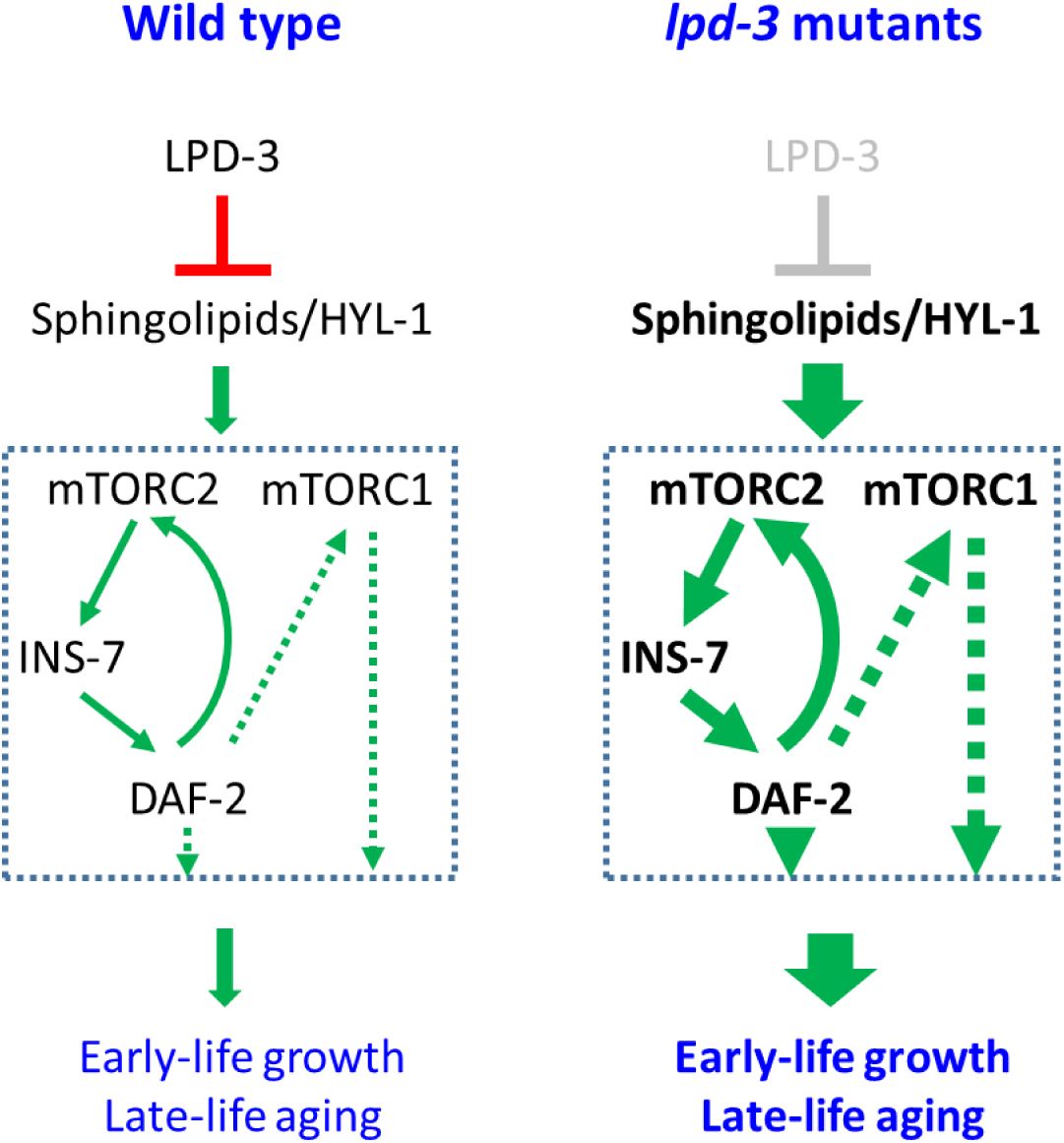
Model of how LPD-3 acts as a brake for aging through sphingolipid and insulin-mTOR regulatory pathways. In the wild type, LPD-3 mediates phospholipid trafficking from ER to PM, and in turn, suppresses sphingolipid levels. Insulin and mTOR signaling are activated at moderate levels to promote early-life growth and late-life aging. In the *lpd-3* mutants, reduced LPD-3 function leads to sphingolipid (hexaceramide type) up-regulation and hyperactivation of insulin and mTOR signaling. INS-7/DAF-2/mTORC2 forms a proposed vicious cycle to sustain insulin and mTOR activation through a positive feedback loop, culminating in hastened late-life aging and shortened lifespan. Red indicates inhibition, while green indicates activation. Solid arrows indicate proposed action based on this study, while dashed arrows indicate regulation inferred from the literature.

**Supplementary Table 1 Lipidomic profiling of phospholipids, glycerolipids and sphingolipids by LC-MS/MS.** 272 mass pairs were monitored by multiple reaction monitoring (MRM) mode. Data were background corrected and normalized with internal standards. Each group had three replicates (n=3) and their respective average (ave), standard deviation (sd), and relative standard deviation (RSD (%)) were calculated. The data file is organized by lipid class/subclass and when applied, their respective fatty acid composition (FA); Triacylglycerides (TAG), hexaceramides (Hexcer), phosphatidylcholine (PC), phosphatidylglycerol (PG), phosphatidylserine (PS), phosphatidylethanolamine (PE), phosphatidylinositol (PI), *lyso*-phosphatidylethanolamine (LPE).

